# Fingerprints of cancer by persistent homology

**DOI:** 10.1101/777169

**Authors:** A. Carpio, L. L. Bonilla, J. C. Mathews, A. R. Tannenbaum

## Abstract

We have carried out a topological data analysis of gene expressions for different databases based on the Fermat distance between the *z* scores of different tissue samples. There is a critical value of the filtration parameter at which all clusters collapse in a single one. This critical value for healthy samples is gapless and smaller than that for cancerous ones. After collapse in a single cluster, topological holes persist for larger filtration parameter values in cancerous samples. Barcodes, persistence diagrams and Betti numbers as functions of the filtration parameter are different for different types of cancer and constitute fingerprints thereof.

---

Ongoing discussions about the causes of cancer include somatic mutations (with related searches to identify relevant molecular pathways in cellular signaling that are cancer drivers) [1], and epigenetics, i.e., heritable changes in gene expression or silencing that do not involve changes to the underlying DNA sequence (phenotype changes without a change in genotype), which in turn affects how cells read the genes [2]. Epigenetic information is controlled by genome sequence, environmental exposure, and stochasticity. It determines development, tissue differentiation, and cellular responsiveness [3]. In Waddington’s landscape [4, 5], a pluripotent stem cell acquires differentiated properties as it falls through a morphogenetic field towards different attractors of the gene regulatory network [3]. Sometimes, specific and anomalous cancer attractors occurring in the epigenetic landscape are postulated and sought [6]. The environment has a profound effect on developmental plasticity, particularly with aging and susceptibility to common disease. Modifications of DNA or associated factors that have information content, other than the DNA sequence itself, are maintained during cell division, influenced by the environment, and cause stable changes in gene expression. Thus, the epigenetic landscape itself evolves dynamically [3]. Epigenetic energy landscapes have been built by viewing methylation maintenance as a communications system and using information theoretic techniques [7, 8]. The duality between the fate of individual cells developing stochastically in Waddington’s landscape of gene expression and the evolution of their density is an important problem studied using optimal transport in Ref. [9].

Be the proximate or ultimate causes of cancer what they may be, differences between healthy people and cancer patients ought to give useful clues. In healthy people, the expressions of many genes are connected in feedback loops that influence and settle cell phenotypes and functions. In cancerous DNA methylation, genes are activated or silenced in ways that differ from healthy methylation. Cancerous cells undergo mutations or change their gene expression. They either lose sufficient feedback loops that leave a smaller set of genes connected or acquire different feedback loops. It is important to characterize the expression values of such genes as they may act as fingerprints for different types of cancer. On the other hand, patients having the same type of cancer should provide evidence of gene mechanisms involved in their disease. Given appropriate sets of related cancer patients, analysis of their gene data should produce the sought evidence. Different techniques of data analysis are important to extract information from high dimensional databases about types of cancer [10–12], cancer response to drugs [13], etc. In Biomedicine, topological data analysis (TDA) has been used for gene expression profiling on breast tumors [10], identification of different types of diabetes [14], spinal cord and brain injury [15], organization of brain activity [16], human recombination [17], cellular differentiation and development [18], or in protein interaction networks [19].

For sets of cancer patients, we utilize a peculiar distance between genes, the *Fermat distance*, which captures relations between genes that are important in different cancer types, and a methodology to distinguish genes based on TDA. The aim is to produce fingerprints of different cancers by analyzing appropriate databases.

Firstly, we reduce the dimension of our data by considering only those genes participating in well-known pathways that appear in the Kyoto Encyclopedia of Genes and Genomes (KEGG, https://www.genome.jp/kegg/). We consider mRNA expression values (transcript count estimates) for 77 genes in the cell cycle gene/protein interaction network for pancreatic adenocarcinoma. These genes are expressed in 177 samples of pancreatic adenocarcinomas in The Cancer Genome Atlas (TCGA in cBioPortal, http://www.cbioportal.org/) [20, 21]. They are also expressed in 248 healthy pancreas samples from GTEx, the Gene Tissue Expression project (https://gtexportal.org). In the two spaces of healthy and cancerous samples, we form point clouds of 77 gene expressions given by *z* scores.

Within the manifolds of gene expression values in the spaces of healthy and cancerous samples, we define a Fermat distance as the shortest geodetic distance between two points divided by an appropriate power of the local density of points. Recall that each point in the set of healthy samples and each point in the set of cancerous samples comprises the value of a single gene expression for all the selected genes. The genes are the same for each set of samples. To compute the Fermat distance between two points (gene expressions), we need to consider all paths between them that comprise intermediate points in the cloud, divide the Euclidean norm of each segment of the path by some power of the local density, add the resulting numbers, and find the Fermat distance as the smallest such number. Note that the Fermat distance is independent of the gene enumeration and considers all the values of the gene expressions for each sample in the two different sets of tissues. *z* scores of gene expressions that are close in Fermat distance are those whose deviations from their mean value are similar when the local density of the point cloud is taken into consideration. This leads to groups of samples more or less deviated from the mean value of gene expression. As the power of the local density increases, the separation between highly deviated and lowly deviated gene expressions becomes larger. In turn, these groupings show how distorted gene expression and gene connections are.

Secondly, we study clusters based on the Fermat distance using TDA. These clusters make noticeable the relations between gene expressions in healthy samples and those in cancerous samples. Fig. 1 compares H_0_ (clusters), H_1_ (cycles) and H_2_ (holes) homologies for healthy and cancerous pancreas samples. As the filtration parameter *r* increases past a critical value *r*_*c*_, samples collapse in a large single cluster. For healthy samples, *r*_*c*_ is smaller than the corresponding value for cancerous samples. While cycles and holes disappear before all clusters form a single one in healthy tissue at *r* = *r*_*c*_, they persist after the formation of a single cluster for cancerous tissue. The Betti number b_0_, which measures the number of independent clusters as a function of *r*, drops faster as *r* increases for healthy tissue. Thus, *r*_*c*_ is smaller for healthy tissue, and the Betti number b_1_ (cycles) vanishes before *r* reaches *r*_*c*_. However, b_1_ *>* 0 persists for cancerous tissue as *r* increases past the value *r*_*c*_. As shown in the example of Fig. 2, this indicates that the gene expressions form irregular clusters with persistent cycles and holes for cancerous tissue, whereas the clusters are more compact for healthy tissue. Fig. 2 gives an idea of the distortion in healthy gene function caused by cancer. Note that the appearance and annihilation of holes give an idea of how clusters begin to link together at certain scales and how compact the larger clusters become. Furthermore, Fig. 1 shows that the higher homology classes H_2_ (holes) appear for cancerous data but are barely noticeable for healthy ones. The same occurs for H_3_, which is not shown in Figs. 1(a) and (c). This gives an idea of the distortion in healthy gene function caused by cancer.

**FIG. 1.**
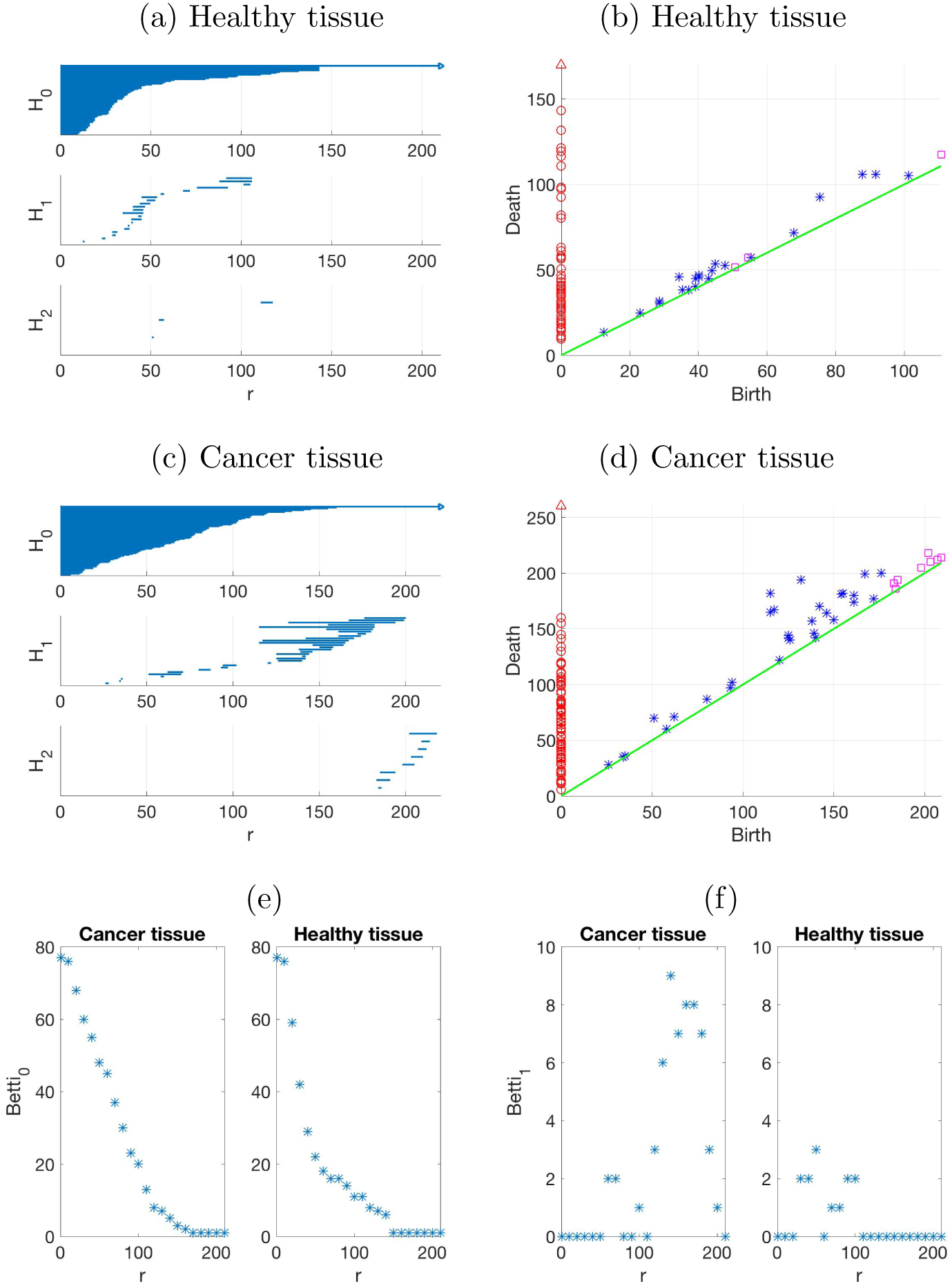
Persistence homology for RNA expression values of 77 genes appearing in the KEGG cell cycle gene/protein interaction network for 177 pancreatic adenocarcinoma tissue samples (TCGA database), and 248 healthy pancreas samples (GTEx project). (a) Barcodes and (b) persistence diagram (healthy tissue), (c) barcodes and (d) persistence diagram (cancer samples), with H_0_ (red circles), H_1_ (blue asterisks), H_2_ (magenta squares). Scales in (b) and (d) are different. (e) b_0_ and (f) b_1_ Betti numbers from Vietoris-Rips filtrations based on Fermat distances.

**FIG. 2.**
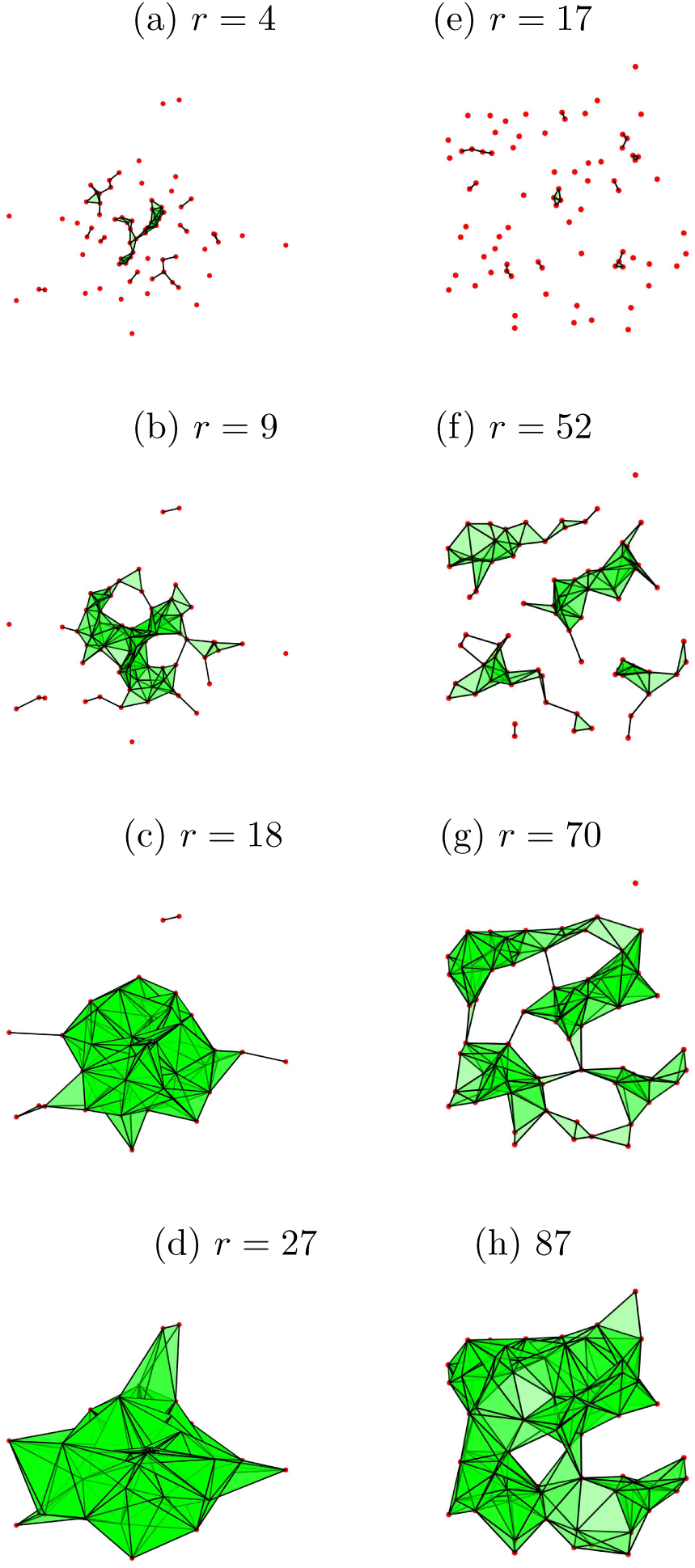
Two examples of bidimensional point clouds (red dots) and their simplicial complexes from the Vietoris-Rips filtration with parameter *r*. (a)-(d) are similar to the homologies H_0_, H_1_ of Fig. 1(a) for healthy tissues, whereas (e)-(h) are similar to cancer tissue homologies in Fig. 1(c).

Thirdly, we perform a similar analysis based on the Fermat distance and TDA for different types of cancer. We analyze how mRNA is expressed in 177 samples of pancreatic adenocarcinoma, 817 samples of breast invasive carcinoma, and 253 sarcoma samples (all data obtained from TCGA lists) for 280 genes listed in the cBioPortal for Cancer Genomics. As shown in Fig. 3, all cancerous samples display holes for some interval beyond the critical value of the filtration parameter *r*_*c*_ for which there is a single cluster. However, the evolution of their Betti numbers with *r*, depicted in Fig. 4, distinguishes different cancer types. The critical *r*_*c*_ is smaller for pancreatic adenocarcinoma, much larger for breast carcinoma and intermediate for sarcoma. The number of cycles measured by b_1_ is similar for pancreatic and breast cancer (about 40) but it is much larger for sarcoma (about a hundred). The sarcoma tag really encompasses different types of cancer and TDA captures such heterogeneity by yielding a much larger number of cycles b_1_ than the other cancer types.

**FIG. 3.**
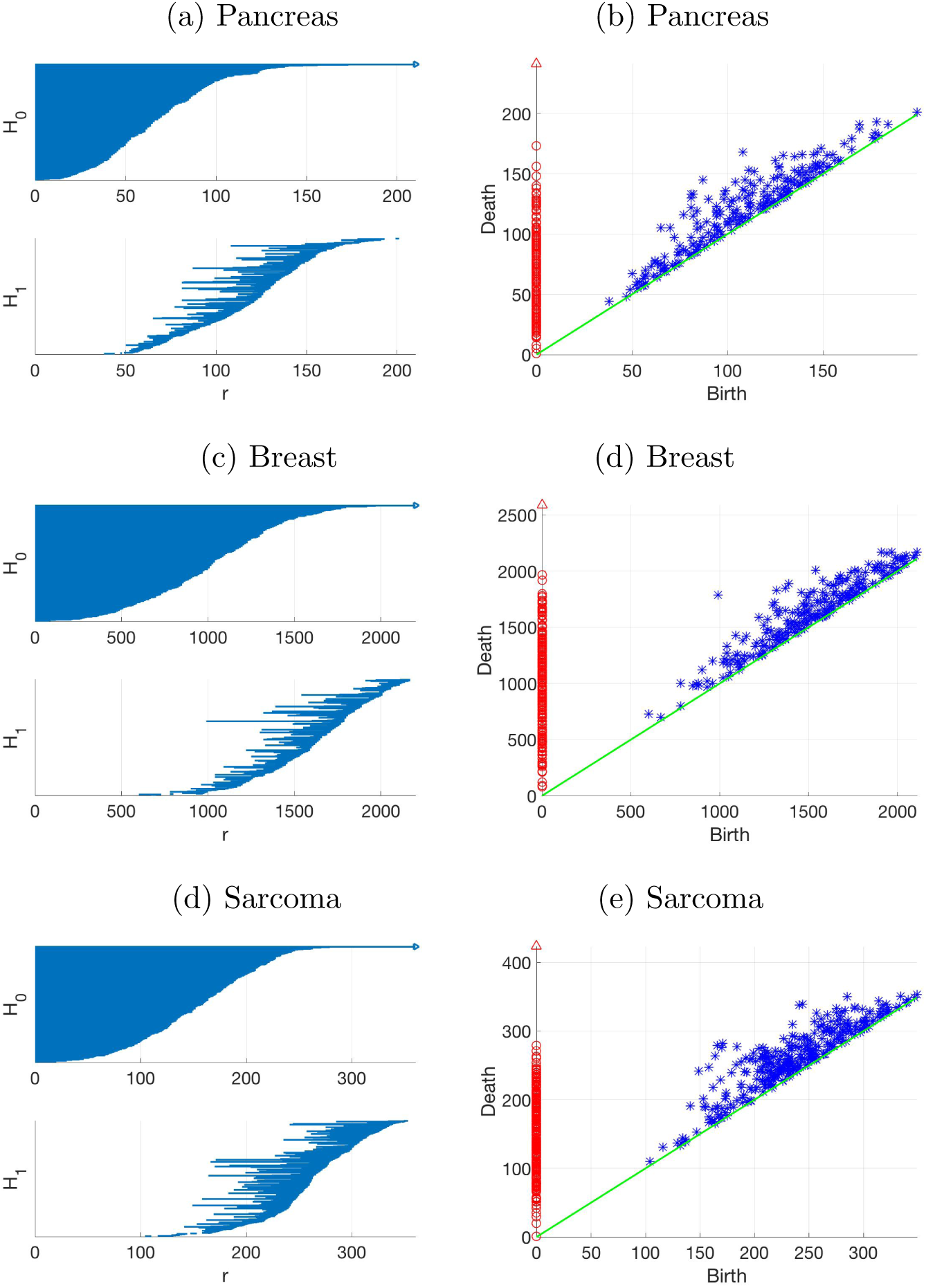
Barcodes and persistence diagrams (H_0_: red circles, H_1_: blue asterisks) for: (a)-(b) 177 pancreatic adenocarcinoma samples, (c)-(d) 817 breast invasive carcinoma samples, (e)-(f) 253 sarcoma samples taken from TCGA. They correspond to mRNA expression values of 280 different genes (cBioPortal for Cancer Genomics) involved in: cell cycle control (34), p53 signaling (6), Notch signaling (55), DNA damage response (12), growth/proliferation signaling (11), survival/cell death regulation signaling (23), telomere maintenance (2), RTK signaling family (16), PI3K-AKT-mTOR signaling (17), Ras-Raf-MEK-Erk/JNK signaling (26), regulation of ribosomal protein synthesis and cellular growth (9), angiogenesis (6), folate transport (5), invasion and metastasis (27), TGF-*β* pathway (43).

**FIG. 4.**
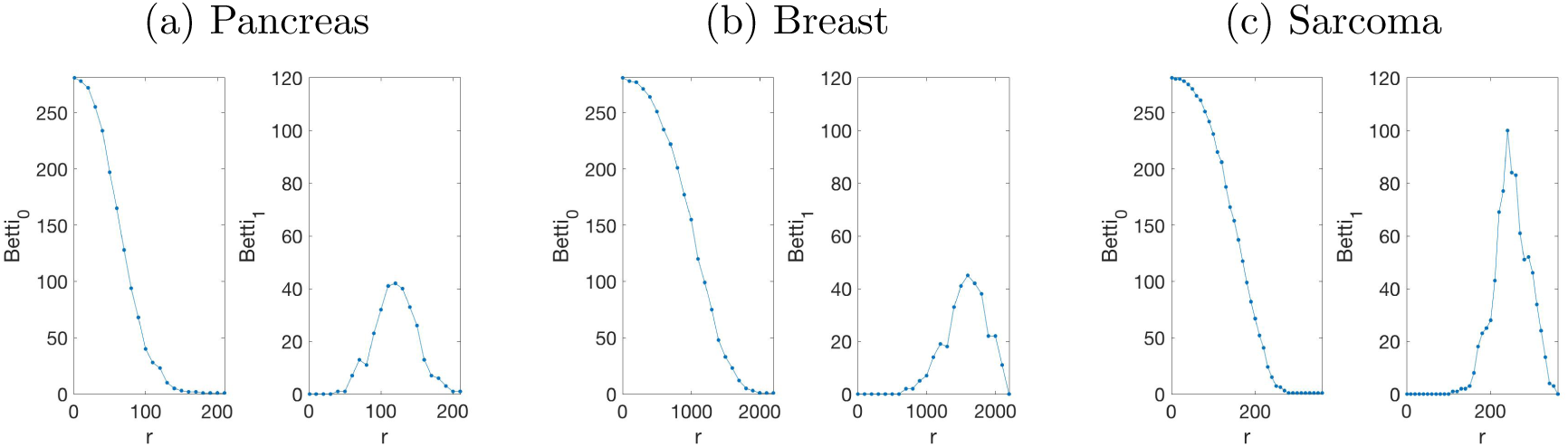
Betti numbers versus filtration parameter *r* for: (a) pancreatic adenocarcinoma, (b) breast invasive carcinoma, (c) sarcoma.

In conclusion, we have carried out a topological data analysis of gene expression values based on the Fermat distance between *z* scores obtained from samples in different databases. The Fermat distance between two gene expressions considers all possible paths joining them in the space of samples in a database, weighs them according to the value of the local point density and finds the minimal value. Thus, this distance provides an average image of the manifolds of gene expressions in databases corresponding to different types of tissue samples. Topological data analysis shows that there is a critical value of the filtration parameter *r*_*c*_ for which all clusters of gene expression values collapse in a single one. This critical value is smaller for healthy samples than for cancerous ones. More importantly, at *r*_*c*_, the collapsed single cluster is gapless for healthy samples, i.e., topological holes have already disappeared, whereas they persist for cancerous samples. Secondly, barcodes, persistence diagrams and Betti numbers as functions of the filtration parameter are different for different types of cancer. These features constitute their fingerprints. There are important open problems to be tackled in the future if we want to use our methodology for personalized medicine. Among them, how to compare new data for single patients with aggregated data and uncertainty quantification.

## I. MATERIALS AND METHODS

### 1. Fermat distance

The Fermat geodetic distance [22] is a distance over geodesics of the manifold *ℳ* that are weighted by the manifold density:

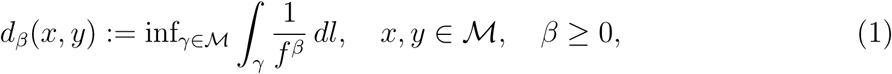

where *γ* are rectifiable curves that start at *x* and end at *y*, and *l*(*x, y*) = *‖x − y‖*_2_ is the Euclidean distance. The minimizer of Eq. (1) privileges trajectories over high density regions and is similar to Fermat’s principle in geometric optics with index of refraction 1*/f* ^*β*^. See Fig. 5. For *β* = 0, Eq. (1) is the isometrical mapping distance (isomap) based on geodesics and Euclidean distances [23]. The variation of the exponent *β* plays a role similar to the variation of the exponent in the multifractal dimension of a chaotic attractor. In the case of the latter, the variation of the exponent distinguishes regions of the attractor that a trajectory visits more often from those scarcely visited [24].

**FIG. 5.**
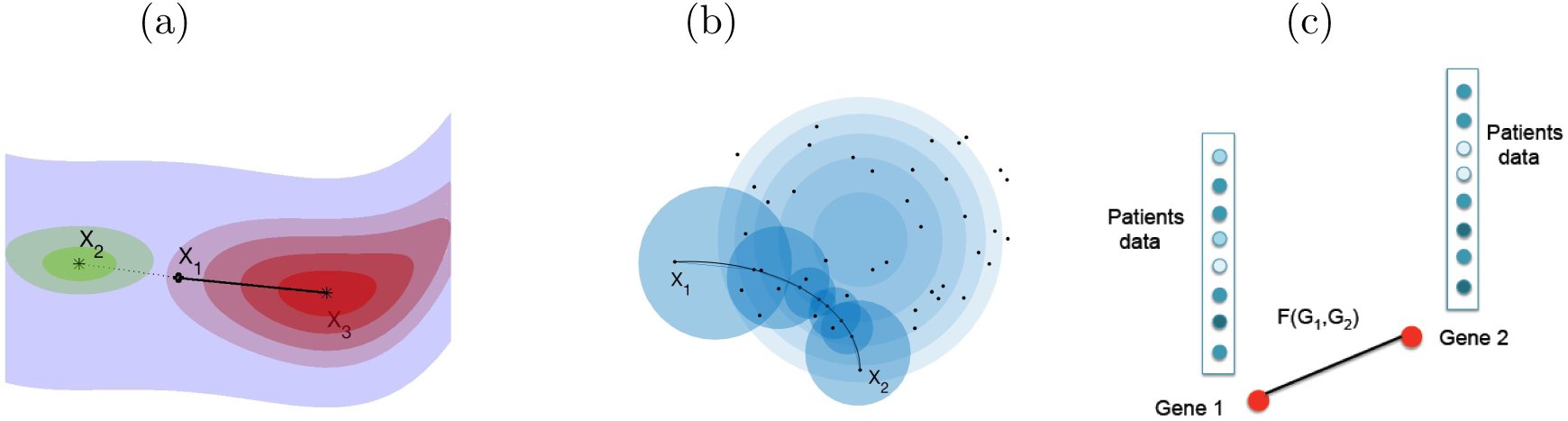
(a) Fermat geodetic distance: *d*_*β*_(*X*_1_, *X*_3_) *< d*_*β*_(*X*_1_, *X*_2_) but, for the Euclidean distance, *‖X*_1_ *− X*_2_*‖*_2_ *< ‖X*_1_ *− X*_3_*‖*_2_; (b) Illustration of the sample Fermat distance *d*_𝕏_*n,α*(*X*_1_, *X*_2_). (c) Measurements of gene expression counts (e.g., mRNA) for a population of patients can be compared using Fermat distances *F* (*G*_1_, *G*_2_), which weight the information provided by all the patients.

In practice, the manifold *ℳ* and its dimension *d* are not known, as we only have a cloud of data points in a space of much higher dimension *D*. To estimate the Fermat distance between points *p* and *q*, we consider 𝕏_*n*_ samples of the manifold *ℳ* with common density *f*, and let *y*_*j*_ *∈* 𝕏_*n*_, *k ≥* 1, be points with *y*_1_, *y*_*k*_ closest to the initial and final points *p, q*, respectively. As the number of samples *n → ∞*, the sample Fermat distance is

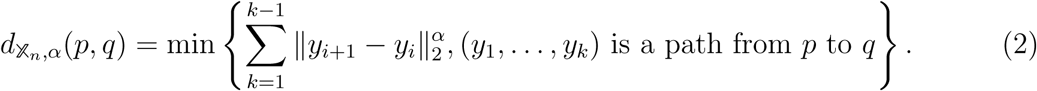

Groisman *et al* have proved that the rescaled sample distance converges to the Fermat distance:

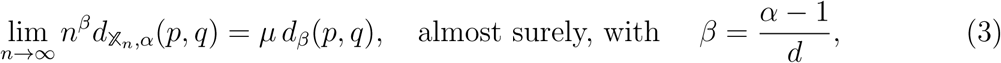

where *µ* is some constant, cf Ref. [25] for the precise statements. In our analysis of databases, we use the sample Fermat distance with variable exponent *α*. We consider the same number of gene expression values for each database of dimension *n* (*n* equals the number of tissue samples in the database, it may differ from one database to another, and it is sufficiently large). The point clouds for gene expression values obtained from healthy and from cancerous tissues provide average images of the corresponding manifolds. Gene *z* scores of *n* samples that are close have a similar deviation from their mean value. As *α* increases above that of the Euclidean distance, *α* = 1, the separation between *z* scores close to the mean and deviated *z* scores becomes sharper. Values between 3 and 6 give good separations and we have typically used *α* = 3 in our computations.

### 2. Persistence homology

Homology distinguishes topological spaces (e.g., circle, annulus, sphere, torus, etc) by quantifying their connected components, topological circles, trapped volumes, and so forth. Suppose we have a finite set of data points from a sampling of the underlying topological space. We measure data homology by creating connections between nearby data points, varying the scale over which these connections are made (= *filtration parameter*), and looking for features that persist across scales [26, 27]. This can be achieved by building the *Vietoris-Rips complex* from all pairwise distances between points in the dataset. For each value of the filtration parameter *r >* 0, we form a simplicial complex *S*_*r*_ by finding all gatherings of *k* + 1 points such that all pairwise distances between these points are smaller than *r*. Each such gathering is a *k*-simplex. The simplicial complex *S*_*r*_ comprises finitely many simplices such that (i) every nonempty subset of a simplex in *S*_*r*_ is also in *S*_*r*_, and (ii) two *k*-simplices in *S*_*r*_ are either disjoint or intersect in a lower dimensional simplex. In *S*_*r*_, 0-simplices are the data points, 1-simplices are edges, connections between two data points, 2-simplices are triangles formed by joining 3 data points through their edges, 3-simplices are tetrahedra, etc. See the example in Fig. 2.

Within the set of all *k*-simplices in *S*_*r*_, we can distinguish closed submanifolds called *k*-cycles, and cycles called boundaries because they are also the boundary of a submanifold. A homology class is an equivalence class of cycles modulo boundaries. A homology class H_*k*_ is the set of independent topological holes of dimension *k*, represented by cycles which are not the boundary of any submanifold. The dimension of H_*k*_ is the *k*th *Betti number* b_*k*_. For instance, b_0_ is the number of connected components, b_1_ is the number of topological circles, b_2_ is the number of trapped volumes, and so on. See Refs. [27, 28] for precise definitions.

Graphical representations of homologies are barcodes and persistence diagrams as in Figures 1 and 3. *Barcodes* are segments of the filtration parameter *r* within homologies that mark the beginning and end of each element thereof. Simplices of higher dimension than vertices are created at some value *r*_*b*_ and disappear at some *r*_*d*_ *> r*_*b*_. For equally spaced values of *r*, we represent each bar in the barcode by a point (*r*_*b*_, *r*_*d*_) in the Cartesian plane, which produces a *persistence diagram*. A point (*x, y*) of the persistence diagram with multiplicity *m* represents *m* features that all appear for the first time at scale *x* and disappear at scale *y* The height of a point over the diagonal, (*y − x*), gives the length of the corresponding bar in the barcode and is called the *persistence* of the feature. Points near the diagonal are inferred to be noise while points further from the diagonal are considered topological signal [26]. Coloring differently different homologies (H_0_, H_1_, etc) we can accumulate plenty of topological information in one 2D persistence diagram, cf Figs. 1 and 3.

### 3. Data

- The 77 genes to compare sick/healthy pancreas tissue are: ABL1,Abl ANAPC10,APC/C ATM,ATM/ATR BUB1,Bub1 BUB1B,BubR1 BUB3,Bub3 CCNA2,CycA CCNB3,CycB CCND1,CycD CCNE1,CycE CCNH,CycH CDC14B,Cdc14 CDC20,Cdc20 CDC25A,Cdc25A CDC25B,Cdc25B CDC45,Cdc45 CDC6,Cdc6 CDC7,Cdc7 CDK1,CDK1 CDK2,CDK2 CDK4,CDK4 CDK7,CDK7 CDKN1A,Cip1 CDKN1B,Kip1 CDKN2A,ARF CDKN2B,Ink4B CDKN2C,Ink4C CDKN2D,Ink4D CHEK1,Chk1 CREBBP,p300 DBF4,Dbf4 E2F1,E2F1 E2F4,E2F4 ESPL1,Esp1 FZR1,Cdh1 GADD45G,GADD45 GSK3B,GSK3B HDAC1,HDAC MAD1L1,Mad1 MAD2L2,Mad2 MCM2,Mcm2 MCM3,Mcm3 MCM4,Mcm4 MCM5,Mcm5 MCM6,Mcm6 MCM7,Mcm7 MDM2,Mdm2 MYC,c-Myc ORC1,Orc1 ORC2,Orc2 ORC3,Orc3 ORC4,Orc4 ORC5,Orc5 ORC6,Orc6 PCNA,PCNA PKMYT1,Myt1 PLK1,Plk1 PRKDC,DNA-PK PTTG2,PTTG RAD21,Rad21 RB1,Rb RBL1,p107 SFN,14-3-3s SKP1,Skp1 SKP2,Skp2 SMAD2,Smad2 SMAD4,Smad4 SMC1B,Smc1 SMC3,Smc3 STAG1,Stag1 TFDP1,Dp-1 TGFB1,TGFB TP53,p53 TTK,Mps1 WEE2,Wee YWHAQ,14-3-3 ZBTB17,Miz1 Here the healthy pancreas samples are from GTEx, the Gene Tissue Expression project (https://gtexportal.org), cancerous samples are from TCGA, PanCancer Atlas in cBio-Portal (http://www.cbioportal.org/) [20, 21].
- Genes and cycles from TCGA, PanCancer Atlas in cBioPortal, http://www.cbioportal.org/) [20, 21]: Cell Cycle control (34): RB1, RBL1, RBL2, CCNA1, CCNB1, CDK1, CCNE1, CDK2, CDC25A CCND1, CDK4, CDK6, CCND2, CDKN2A, CDKN2B, MYC, CDKN1A, CDKN1B, E2F1, E2F2, E2F3, E2F4, E2F5, E2F6, E2F7, E2F8, SRC, JAK1, JAK2, STAT1, STAT2, STAT3, STAT5A, STAT5B. p53 signaling (6): TP53, MDM2, MDM4, CDKN2A, CDKN2B, TP53BP1. Notch signaling (55): ADAM10, ADAM17, APH1A, APH1B, ARRDC1, CIR1, CTBP1, CTBP2, CUL1, DLL1, DLL3, DLL4, DTX1, DTX2, DTX3, DTX3L, DTX4, EP300, FBXW7, HDAC1, HDAC2, HES1, HES5, HEYL, ITCH, JAG1, JAG2, KDM5A, LFNG, MAML1, MAML2, MAML3, MFNG, NCOR2, NCSTN, NOTCH1, NOTCH2, NOTCH3, NOTCH4, NRARP, NUMB, NUMBL, PSEN1, PSEN2, PSENEN, RBPJ, RBPJL, RFNG, SNW1, SPEN, HES2, HES4, HES7, HEY1, HEY2. DNA damage (12): CHEK1, CHEK2, RAD51, BRCA1, BRCA2, MLH1, MSH2, ATM, ATR, MDC1, PARP1, FANCF. Proliferation (11): CSF1, CSF1R, IGF1, IGF1R, FGF1, FGFR1, AURKA, DLEC1, PLAGL1, OPCML, DPH1. Survival/death cell (23): NFKB1, NFKB2, CHUK, DIRAS3, FAS, HLA-G, BAD, BCL2, BCL2L1, APAF1, CASP9, CASP8, CASP10, CASP3, CASP6, CASP7, GSK3B, ARL11, WWOX, PEG3, TGFB1, TGFBR1, TGFBR2. Telomere maintenance (2): TERC, TERT. RTK signaling (16): EGFR, ERBB2, ERBB3, ERBB4, PDGFA, PDGFB, PDGFRA, PDGFRB, KIT, FGF1, FGFR1, IGF1, IGF1R, VEGFA, VEGFB, KDR. PI3K-AKT-mTOR signaling (17): PIK3CA, PIK3R1, PIK3R2, PTEN, PDPK1, AKT1, AKT2, FOXO1, FOXO3, MTOR, RICTOR, TSC1, TSC2, RHEB, AKT1S1, RPTOR, MLST8. Ras-Raf-MEK-Erk/JNK signaling (26): KRAS, HRAS, BRAF, RAF1, MAP3K1, MAP3K2, MAP3K3, MAP3K4, MAP3K5, MAP2K1, MAP2K2, MAP2K3, MAP2K4, MAP2K5, MAPK1, MAPK3, MAPK4, MAPK6, MAPK7, MAPK8, MAPK9, MAPK12, MAPK14, DAB2, RASSF1, RAB25. Ribosomal protein synthesis and cell growth (9): RPS6KA1, RPS6KA2, RPS6KB1, RPS6KB2, EIF5A2, EIF4E, EIF4EBP1, RPS6, HIF1A. Angiogenesis (6): VEGFA, VEGFB, KDR, CXCL8, CXCR1, CXCR2. Folate transport (5): SLC19A1, FOLR1,, FOLR2, FOLR3, IZUMO1R. Invasion and metastasis (27): MMP1, MMP2, MMP3, MMP7, MMP9, MMP10, MMP11, MMP12, MMP13, MMP14, MMP15, MMP16, MMP17, MMP19, MMP21, MMP23B, MMP24, MMP25, MMP26, MMP27, MMP28, ITGB3, ITGAV, PTK2, CDH1, SPARC, WFDC2. TGF-beta (43): TGFB1, TGFB2, TGFB3, TGFBR1, TGFBR2, TGFBR3, BMP2, BMP3, BMP4, BMP5, BMP6, BMP7, GDF2, BMP10, BMP15, BMPR1A, BMPR1B, BMPR2, ACVR1, ACVR1B, ACVR1C, ACVR2A, ACVR2B, ACVRL1, Nodal, GDF1, GDF11, INHA, INHBA, INHBB, INHBC, INHBE, SMAD2, SMAD3, SMAD1, SMAD5, SMAD4, SMAD9, SMAD6, SMAD7, SPTBN1, TGFBRAP1, ZFYVE9.

## ACKNOWLEDGMENTS

This work has been supported by the FEDER/Ministerio de Ciencia, Innovación y Universidades – Agencia Estatal de Investigación grants MTM2017-84446-C2-1-R (AC) and MTM2017-84446-C2-2-R (LLB). AC and LLB thank Russel Caflisch for hospitality during a sabbatical stay at the Courant Institute. We thank Joseph O. Deasy, Saad Nadeem, Jung Hun Oh, and Maryam Pouryahya for useful discussions and comments.

